# Identifying locations susceptible to micro-anatomical reentry using a spatial network representation of atrial fibre maps

**DOI:** 10.1101/2021.09.13.460069

**Authors:** Max Falkenberg, James A Coleman, Sam Dobson, David J Hickey, Louie Terrill, Alberto Ciacci, Belvin Thomas, Nicholas S Peters, Arunashis Sau, Fu Siong Ng, Jichao Zhao, Kim Christensen

**Affiliations:** Centre for Complexity Science, Imperial College London, London, United Kingdom; Department of Physics, Imperial College London, London, United Kingdom; ElectroCardioMaths Programme, Imperial Centre for Cardiac Engineering, Faculty of Medicine, Imperial College London, London, United Kingdom; Auckland Bioengineering Institute, The University of Auckland, Auckland, New Zealand

## Abstract

Micro-anatomical reentry has been identified as a potential driver of atrial fibrillation (AF). In this paper, we introduce a novel computational method which aims to identify which atrial regions are most susceptible to micro-reentry. The approach, which considers the structural basis for micro-reentry only, is based on the premise that the accumulation of electrically insulating interstitial fibrosis can be modelled by simulating percolation-like phenomena on spatial networks. Our results suggest that at high coupling, where micro-reentry is rare, the micro-reentrant substrate is highly clustered in areas where the atrial walls are thin and have convex wall morphology. However, as transverse connections between fibres are removed, mimicking the accumulation of interstitial fibrosis, the substrate becomes less spatially clustered, and the bias to forming in thin, convex regions of the atria is reduced. Comparing our algorithm on image-based models with and without atrial fibre structure, we find that strong longitudinal fibre coupling can suppress the micro-reentrant substrate, whereas regions with disordered fibre orientations have an enhanced risk of micro-reentry. We suggest that with further development, these methods may have future potential for patient-specific risk stratification, taking a longitudinal view of the development of the micro-reentrant substrate.

**Author summary:** Atrial fibrillation (AF) is the most common abnormal heart rhythm, yet, despite extensive research, treatment success rates remain poor. In part, this is because there is an incomplete understanding of the mechanistic origin of AF. In this paper, we investigate one proposed mechanism of AF, the formation of “micro-reentrant circuits”, which can be thought of as a “short circuit”, forming when electrically insulating fibrosis (structural repair tissue) infiltrates the space between heart muscle cells. Previously, such circuits have been found in experimental hearts, but identifying these circuits clinically is difficult. Here, we aim to take a small step towards developing computational methods for identifying where in the atria these circuits are most likely to form, drawing on techniques from network science. Our approach indicates that a number of factors are key to determining where circuits form, most notably the thickness of the heart muscle, and the alignment of muscle fibres.

## 1 Introduction

Atrial fibrillation (AF) is the most common cardiac arrhythmia with significant impacts on both morbidity and mortality [1]. Despite extensive research, there are significant disagreements within the cardiac electrophysiology community as to the mechanisms underlying AF [2, 3].

Most studies into the mechanistic origin of AF focus on the maintenance of AF, typically arguing for either organised (mother waves or stable rotors) or disorganised mechanisms (multiple reentrant wavelets). However, if both organised and disorganised mechanisms of AF coexist on a continuous spectrum of electromechanical organisation, as suggested recently [4], it is unlikely that any single treatment strategy will be successful across the full spectrum of AF mechanisms. It is for this reason that personalised, patient specific approaches to AF treatment have become a key research focus in recent years [5, 6].

Here we computationally investigate one mechanism of AF initiation and maintenance, the formation of micro-anatomical reentrant circuits; continuously activated electrical circuits anchored to the fibre structure of the atria [7]. Importantly, the size of such circuits is often at, or below, the spatial resolution which can be resolved with conventional multi-electrode mapping [7, 8]. Similarly, the interstitial fibrosis which insulates these circuits is often not easy to detect using conventional LGE-MRI [4], and extracting precise fibrosis densities is challenging given the variability in signal thresholding choices between scans and patients [9]. Hence, computational approaches allow for a degree of hypothesis testing which avoids some of these challenges.

In this proof of concept, we introduce a novel method to assess the feasibility of patient-specific predictions for the distribution of micro-anatomical reentry across the atria. In particular, the method is inspired by the idea that it may be possible to predict the emergence of micro-reentry before the atria have accumulated sufficient interstitial fibrosis, in contrast to most other patient specific approaches which take a static view of the AF substrate [5, 10].

Starting from image-based models of the atria, we combine data regarding the atrial geometry and the underlying myocardial fibre structure to form a spatial network. By progressively removing connections in the atrial structure, we assess where in the network a path exists which is sufficiently long to harbour micro-reentry, an approach related to the study of percolation in network science [11]. In the current work, our primary goal is to understand the utility of the method and demonstrate its future potential. With this in mind, our research focuses on how the regions predicted by our method of being susceptible to micro-reentry depend on fibre structure and atrial geometry.

## 2 Aims

It has recently been shown “that human AF may be driven by microanatomic reentrant AF drivers anchored to fibrotically insulated tracks within the complex atrial wall” [8]. To ensure that a given fibre tract can sustain micro-reentry, the reentrant pathway through a fibrotically insulated fibre tract must be at least one refractory wavelength long.

Our approach is illustrated schematically in Fig. 1. Denoting the minimum refractory wavelength during fibrillation as *τ*, our aim is to assess where in the atrial structure fibrotically insulated reentrant pathways exist of length *ℓ*, such that *ℓ* ≥ *τ*. In a healthy heart with minimal interstitial fibrosis, atrial myofibres are well coupled in both the longitudinal and transverse directions. Hence, the longest possible reentrant path is significantly shorter than one refractory wavelength, *ℓ* ≪ *τ*. As the density of electrically insulating interstitial fibrosis increases, the longest reentrant pathway increases until a threshold density is reached at which *ℓ* ≥ *τ* and the fibrotically insulated fibre tract can sustain continuous reentry during AF. Importantly however, this threshold density may vary across different regions of the atria; some regions may be susceptible to micro-reentry at a low fibrosis density, whereas other regions may require a significantly higher fibrosis density before micro-reentry can be induced. If the density of fibrosis is too high in a region, micro-reentry may be prevented due to the absence of any closed loops (compact fibrosis).

**Fig 1.**
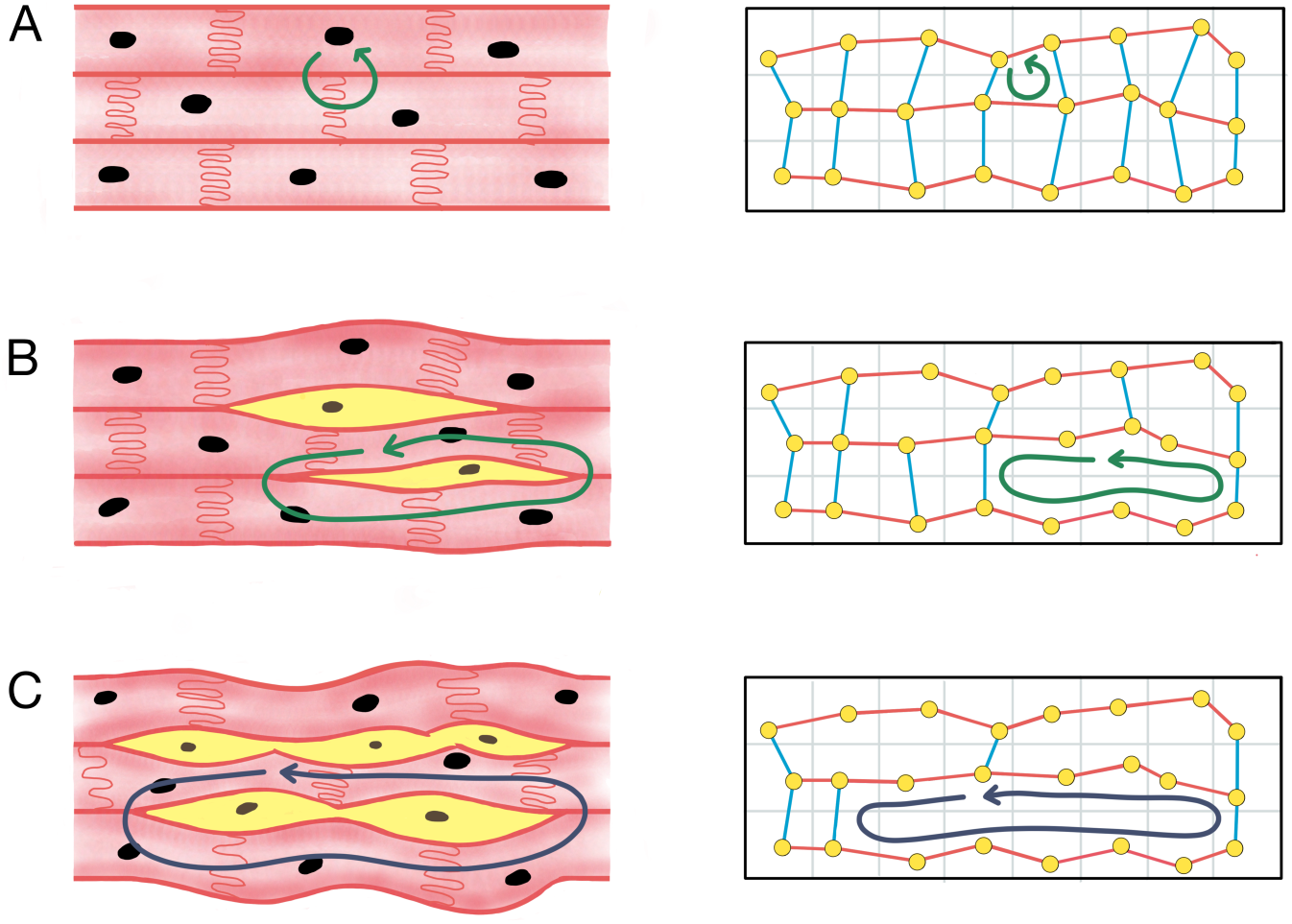
A schematic illustrating our approach. Left column: (A) In the healthy atrial myocardium, cardiomyocytes (pink cells) are well coupled along and across the principle fibre directions, such that the longest re-entrant loop is too short to sustain micro-anatomical reentry (green arrows). (B) Over time, the atria accumulate interstitial fibrosis (yellow cells) forming short segments of electrically isolated fibre. (C) The length of these segments grows as more interstitial fibrosis accumulates until the isolated segments are sufficiently long to harbour micro-reentry (blue arrows). We hypothesise that the accumulation of interstitial fibrosis can be modelled as a spatial network. Right column: Nodes (yellow points) representing a group of cardiomyocytes are connected to their neighbours along fibres (red links) and across fibres (blue links). If the density of interstitial fibrosis is low, there are many transverse (blue) links. We model the increase in the density of interstitial fibrosis as the progressive removal of transverse links, and equate the micro-anatomical reentrant substrate to regions of the spatial network where a loop exists that is longer than one refractory wavelength.

Our hypothesis is that the process of electrically insulating fibre tracts with interstitial fibrosis can be modelled using spatial networks. A spatial network is a graph consisting of nodes representing entities in space, and edges representing spatial connections between those entities [12]. In our case, a node represents one or more atrial cardiomyocytes in a specific atrial region, and an edge between two nodes indicates whether those groups of myocytes are electrically coupled, that is whether an activation wavefront can pass directly from the group of myocytes represented by one node, to the group of myocytes at the other node.

Starting from a network in which the local density of edges is high, our assertion is that the accumulation of insulating interstitial fibrosis in the atria is structurally equivalent to the progressive removal of edges in our spatial network. If the density of fibrosis is low, then the density of edges in the spatial network is high, and similarly, if the density of fibrosis is high, the density of edges in the spatial network is low. Hence, by identifying loops in the spatial network of length *ℓ* ≥ *τ*, we can identify which atrial regions may act as a substrate for micro-anatomical reentry at a given fibrosis density.

## 3 Materials and methods

In this paper, we generate spatial networks from three atrial datasets (see below), each consisting of a voxel mesh with local fibre orientation vectors. For each dataset, we generate spatial networks according to two different approaches, one which preserves the atrial fibre orientations in the spatial network (the fibre model), and a null model approach which ignores the fibre orientation data but retains other structural information such as wall thickness (the fibre-less null model). These two approaches are shown schematically in Fig. 2. By utilising both approaches, we are able to identify how the underlying atrial fibre structure affects the observed micro-reentrant substrate.

**Fig 2.**
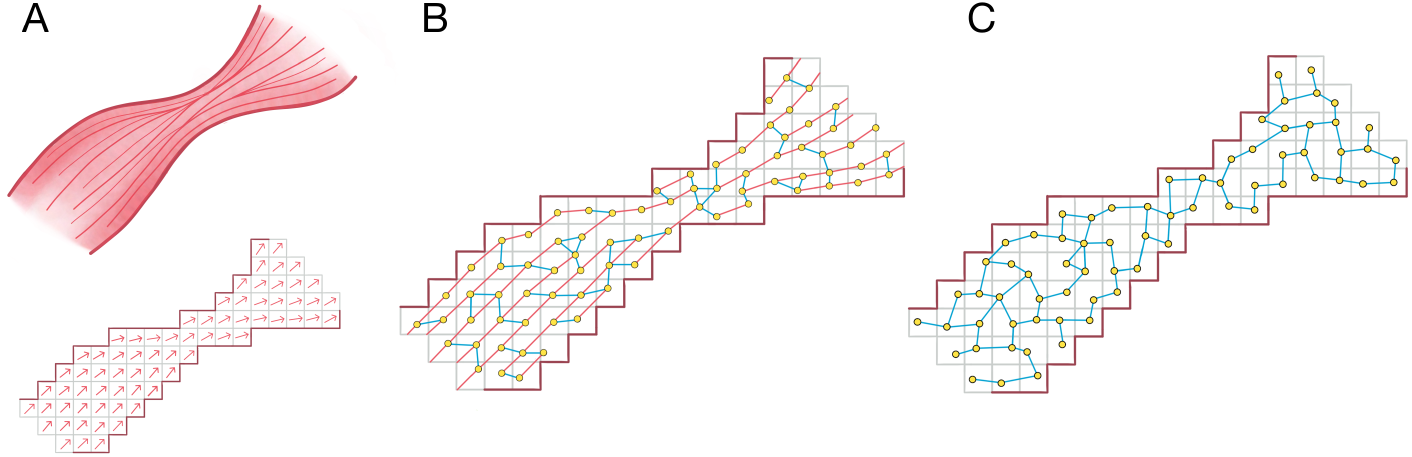
(A) An image based model of the atria is derived from atrial imaging. The model, represented by a mesh of voxels, is coupled with vectors representing the underlying fibre orientation at each point. (B) In the fibre model, a spatial network is generated where nodes (yellow points) are connected with probability 1 along fibre tracts (pink edges), and low probability across fibre tracts (blue edges). (C) In the fibre-less null model, the probability of forming an edge is not biased by the underlying fibre orientation.

### 3.1 Atrial datasets

The spatial network algorithm is tested on three atrial datasets, one human dataset with MRI derived geometry, but synthetic fibre orientations, and two sheep atrial datasets where the fibre structure has been inferred from high resolution serial surface imaging. Each dataset provides a 3d grid of vectors indicating the local fibre orientation at each point in the atria. The sheep datasets are high resolution and inferred directly from the anatomy of each dataset. Summary statistics for the three image-based atrial fibre maps are given in Table 1.

**Table 1.** Summary information for the three atrial fibre orientation datasets.

|  | Human | Sheep<br>Healthy | Sheep<br>Heart failure |
| --- | --- | --- | --- |
| Imaging Method | MRI | Serial Surface Imaging | Serial Surface Imaging |
| Original Image Resolution | $330\mu m$ | $50\mu m$ | $25\mu m$ |
| Fibre Orientation Extraction | Estimated | Anatomically derived | Anatomically derived |
| Fibre Orientation Method | Semi-automated rule based approach focusing on key bundles | Eigen-analysis of structure tensor | Eigen-analysis of structure tensor |
| Resolution after Coarse Graining | $330\mu m$ | $300\mu m$ | $300\mu m$ |
| Dataset Reference | [13, 14] | [15] | [16] |

#### 3.1.1 Synthetic atrial fibre map

The human dataset is detailed in [14] based on techniques developed in [13]. The atrial geometry used provides no information as to the atrial fibre structure. Therefore, a synthetic fibre map is constructed using a semi-automated rule-based approach which is described in the supplementary material (SM) section 1.1.1.

#### 3.1.2 Anatomically derived fibre maps

We study two anatomically derived sheep atrial datasets; one healthy sheep acquired in [15] and a sheep with pacing-induced heart failure [16]. These datasets are individual-specific, retaining significant local heterogeneity in the fibre maps, and are provided at high resolution, in contrast to the synthetic human fibre map. Information regarding the data acquisition process is provided in SM section 1.1.2. In this proof of concept, our aim is not to assess the differences in micro-anatomical reentry between humans and sheep, or between healthy and heart failure (HF) sheep, but to understand how different sources of fibre orientation data effect our results.

### 3.2 Constructing spatial networks with fibre structure

The schematic shown in Fig. 3 outlines the process by which each image-based model is converted into a spatial network in which the underlying fibre structure is retained.

**Fig 3.**
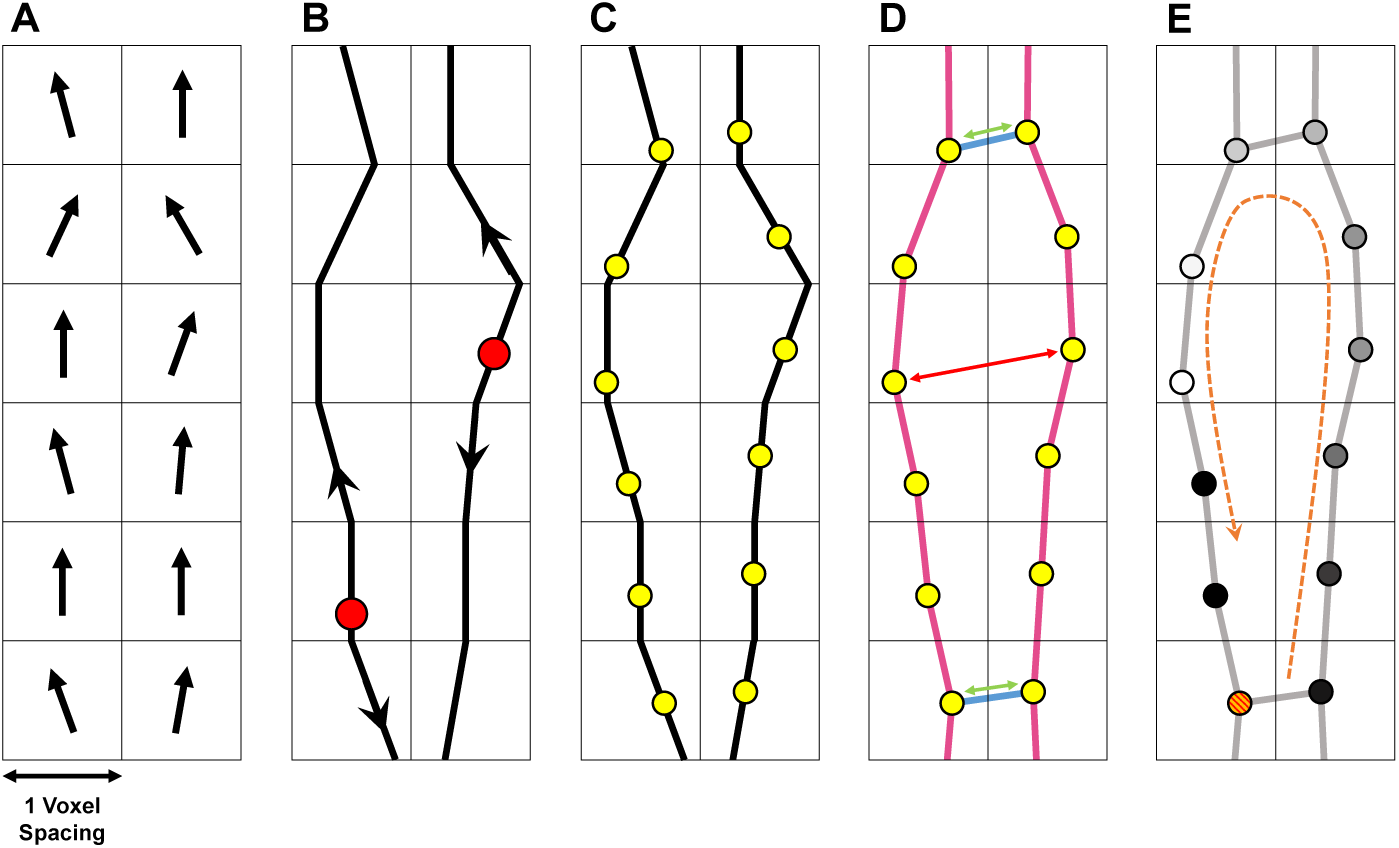
A 2D schematic showing the conversion of local fibre orientation vectors into global fibre tracts which are seeded with nodes and coupled into a spatial network. (A) Each image-based model provides a vector field for the local fibre orientation. (B) Beginning from a set of random seed points (red circles), fibres are generated using tractography, propagating forward and backwards in the direction of the local orientation vector. (C) Nodes (yellow circles) are placed at approximately equal intervals along the fibres. (D) Nodes are coupled to neighbouring nodes with probability 1 if nodes are adjacent and in the same fibre (pink edges), and according to a monotonically decreasing function of distance, *p*(*x*; *r, c*), otherwise (blue edges). Nodes which are close together (green arrows) are more likely to be connected than those which are far apart (red arrows). (E) By applying a discrete diffusion model (see section 3.4) to the network, we identify the micro-reentrant substrate by observing where conduction block (red/orange circle) results in a wavefront (white circle) re-entering a blocked fibre.

#### 3.2.1 Fibre tractography

Fibre tracts are generated across the atrial geometry by applying a modified version of the Evenly Spaced Streamlines (ESS) algorithm, see [17]: (1) Local fibre orientation data for a given atrial dataset is coarse grained to a specific voxel resolution, Fig. 3(A). (2) Placing seed points randomly in the atrial mesh, separated by at least *d*_*sep*_ = 0.7, global fibre tracts are constructed by propagating forwards and backwards along the local fibre orientation vectors until terminated according to a set of pre-specified rules, Fig. 3(B). (3) After fibre tracts have been generated to cover the atrial mesh at an approximately fixed density of 1 node per voxel, points are placed evenly along each fibre, Fig. 3(C). Importantly, unlike other common tractography methods, this modified ESS approach ensures that the density of fibres is approximately uniform across the atrial structure. Technical details for the tractography process are provided in SM section 1.2.

#### 3.2.2 Generation of spatial network structure

The fibres generated from tractography are not connected to each other, Fig. 3(C). To connect fibres in a single spatial network, nodes are placed at even intervals (steps of 1 voxel length) along fibres starting at a random seed point. These nodes connect to neighbouring nodes in the same fibre with probability *p* = 1. Any two nodes which do not lie on the same fibre are connected with probability

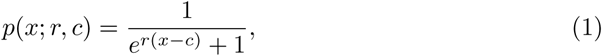

where *x* is the distance between the nodes, *r* = 7 is an arbitrary steepness parameter, and *c* is a characteristic distance. Connecting nodes according to a distance dependent attachment function is a standard technique when studying percolation on spatial networks [11]. For better performance, only nodes separated by *x <* 2 are considered for connection. By varying the spatial coupling parameter *c*, the number of transverse connections between fibres can be controlled from a state where fibres are strongly connected at *c* ≈ 1, to a state where fibres become increasingly disconnected as *c* → 0.

For each dataset, spatial networks are generated with five values of the characteristic distance, *c*. The specific values used and corresponding risk parameters (see section 3.5.1), are given in Table S2 in SM section 1.3.1. For convenience, specific parameter choices are labelled as having a low, medium or high risk of micro-reentry.

### 3.3 Constructing spatial networks without fibre structure

For comparison purposes, we consider a null model where the spatial network excludes fibre orientation. All other geometric information is retained. The fibre-less spatial network is generated analogously to the fibre model, with the omission of the initial fibre tractography steps, illustrated schematically in Fig. 4. Nodes are randomly distributed across each atrial mesh at a density of approximately 1 node per voxel and are added to the spatial network if the distance between the new node and any existing nodes is greater than or equal to *d*_*sep*_ = 0.7; this prevents an over-density of nodes in any one location.

**Fig 4.**
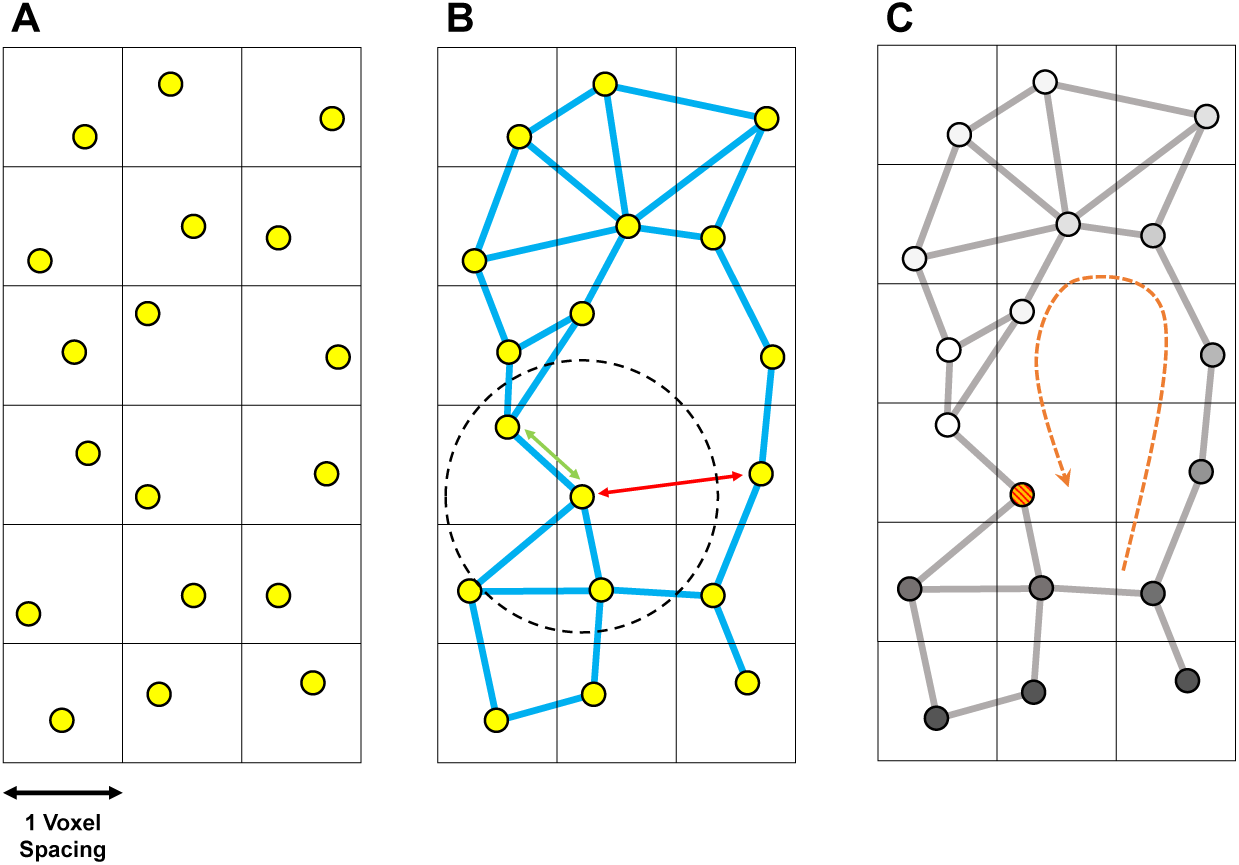
A 2d schematic demonstrating the construction of the spatial network using the fibre-less null model. The process is similar to Fig. 3 where steps (A) and (B) are skipped. (A) Nodes are distributed uniformly in each voxel. (B) Nodes are connected according to a monotonically decreasing function of distance, *p*(*x*; *r, c*), Eq. (1), where *c* is a characteristic distance (dashed circle). There is no directional bias in the connection probability (blue edges only). Nodes which are closer together than the characteristic distance (green arrows) are more likely to be connected than those which are far apart (red arrows). (C) The micro-reentrant substrate is identified using the DDM.

To construct the spatial network, nodes are connected using the same distance dependent probability function as the fibre-model given in Eq. (1). Unlike the fibre-model, none of the nodes are already connected by edges which represent the underlying fibre structure of the atria (no pink edges); the probability of connecting two nodes is independent of their orientation.

For each atrial dataset, we generate spatial networks for three values of the characteristic coupling, corresponding to high, medium and low risk cases, see Table S3 in SM section 1.3.2. The characteristic coupling values for the fibre-less null model are significantly larger than for the fibre model since there are no permanent longitudinal edges.

### 3.4 Identifying the substrate for micro-anatomical reentry

For both the fibre and fibre-less null models, we associate the substrate for micro-anatomical reentry with regions in the corresponding spatial network where closed loops exist of length *ℓ* ≥ *τ*. Here, *τ* represents a typical refractory wavelength. These are found by applying a discrete diffusion model (DDM) to the network, see SM section 1.4. The DDM is related to several extremely simple physics models of micro-anatomical reentry [18–22]. These models are not electrophysiologically realistic models of AF. The strength of discrete models lies in the ease with which structural discontinuities can be modelled, in contrast to conventional models which are significantly complicated when no-flux boundary conditions are imposed at the local level [23]. This justifies the use of the DDM in the current context where we focus exclusively on the structural basis for micro-reentry. However, this approach is not suitable for studying the dynamics of micro-anatomical reentry. For a wider discussion of the role of model choice see [24].

### 3.5 Analysis Methods

#### 3.5.1 Defining risk of micro-anatomical reentry

To derive a measure for the risk of micro-reentry in each spatial network, we use the DDM and assume that all nodes in the network are equally susceptible to conduction block. If each structure has a similar number of nodes which, if blocked, may initiate a micro-anatomical reentry, then the probability of initiating any given circuit is approximately constant. Hence, for a spatial network with *N* potential substrates, the rate, *λ*, at which new micro-reentries are initiated in the DDM is directly related to the number of substrates *N*. This argument follows directly from similar arguments in [20]. A mathematically precise formulation of this argument is provided in SM section 1.5. The rate parameter, *λ*, is used as our measure of the risk of micro-anatomical reentry.

#### 3.5.2 Spatial clustering of reentrant circuits

For each spatial network, we measure how spatially clustered the identified micro-reentrant substrate is. To do so, we sample 1000 circuits from the set of identified risk substrates for each network and extract the coordinate of each detected circuit. The number of independent spatial clusters is then derived by using the DBSCAN algorithm, a standard algorithm for spatial data clustering, see [25]. We set `eps` = 10 (≈ 3*mm*), which is the parameter controlling the distance between points for these to be considered in the same cluster. The number of points required for a cluster is set to 1. If only one cluster is identified by DBSCAN, this indicates that all 1000 sampled circuits in a given network fall within a single, small region of the atrial structure. Conversely, if 1000 clusters are identified, no two circuits are identified in the same location.

#### 3.5.3 Atrial wall thickness & occupied voxel fraction

Wall thickness is a common measurement used when analysing atrial geometry. One issue when calculating wall thickness is that it is not easily defined in the atrial bulk. On the surfaces, wall thickness can be defined as the distance through the atrial wall along the perpendicular surface vector (there are competing, but similar definitions). However, in the bulk many perpendicular surface vectors pass through the same voxel so that the thickness is not uniquely defined.

To resolve this issue, we introduce a novel measure analogous to wall thickness, the occupied voxel fraction (OVF), which quantifies the proximity of voxels to the atrial walls and captures differences in the local wall curvature and thickness gradients, see SM section 1.6. The measure is defined as the number of voxels within a radius 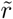 (approximately 1.5mm) which are inside the atrial structure, normalised by the total number of voxels within a sphere of radius 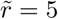. The OVF increases (decreases) if the atrial walls become thicker (thinner), and distinguishes between regions with convex, flat, or concave wall morphologies.

#### 3.5.4 Longitudinal connection fraction

For the spatial networks with fibre structure, we would like to associate the identified micro-reentrant substrate with the degree of local fibre alignment in a particular atrial region. Fibre alignment is not naturally defined in a spatial network. Therefore, we use the longitudinal connection fraction (LCF) as a proxy measure. This is defined as the average number of longitudinal connections (along the fibre direction), normalised by the total number of connections, either longitudinal or transverse. If the LCF in a voxel is large, then fibres in that region are well aligned. Conversely, if the LCF is small, the fibre structure is highly disordered in a local area. More detail is provided in SM section 1.7 where we validate that the LCF in each spatial network accurately reflects the expected degree of fibre misalignment measured in the underlying data.

It is interesting to note that we can compare the LCF to the degree of microscopic anisotropy, calculated using Eq. (1), at the highest level of spatial coupling where micro-anatomical reentry is inducible in our spatial networks. This demonstrates that the onset of micro-reentry in our spatial networks occurs at approximately the same degree of structural anisotropy as found experimentally by Spach *et al*. [26], see SM section 1.8 for details.

## 4 Results

### 4.1 Spatial distribution of identified micro-reentrant substrate

Figures 5 & 6 show the substrate identified as susceptible to micro-reentry in the human atria using the fibre and fibre-less null models respectively, at low, medium and high risk (progressively larger reentry rate, *λ*). Note that low, medium and high risk refers to the overall risk of micro-reentry across the whole atrial tissue, and does not refer to whether specific regions identified are at low or high risk. The spatial distributions of risk for the sheep atria are shown in SM section 2.1. How the differences in our datasets effect our results is discussed in SM section 2.2.

**Fig 5.**
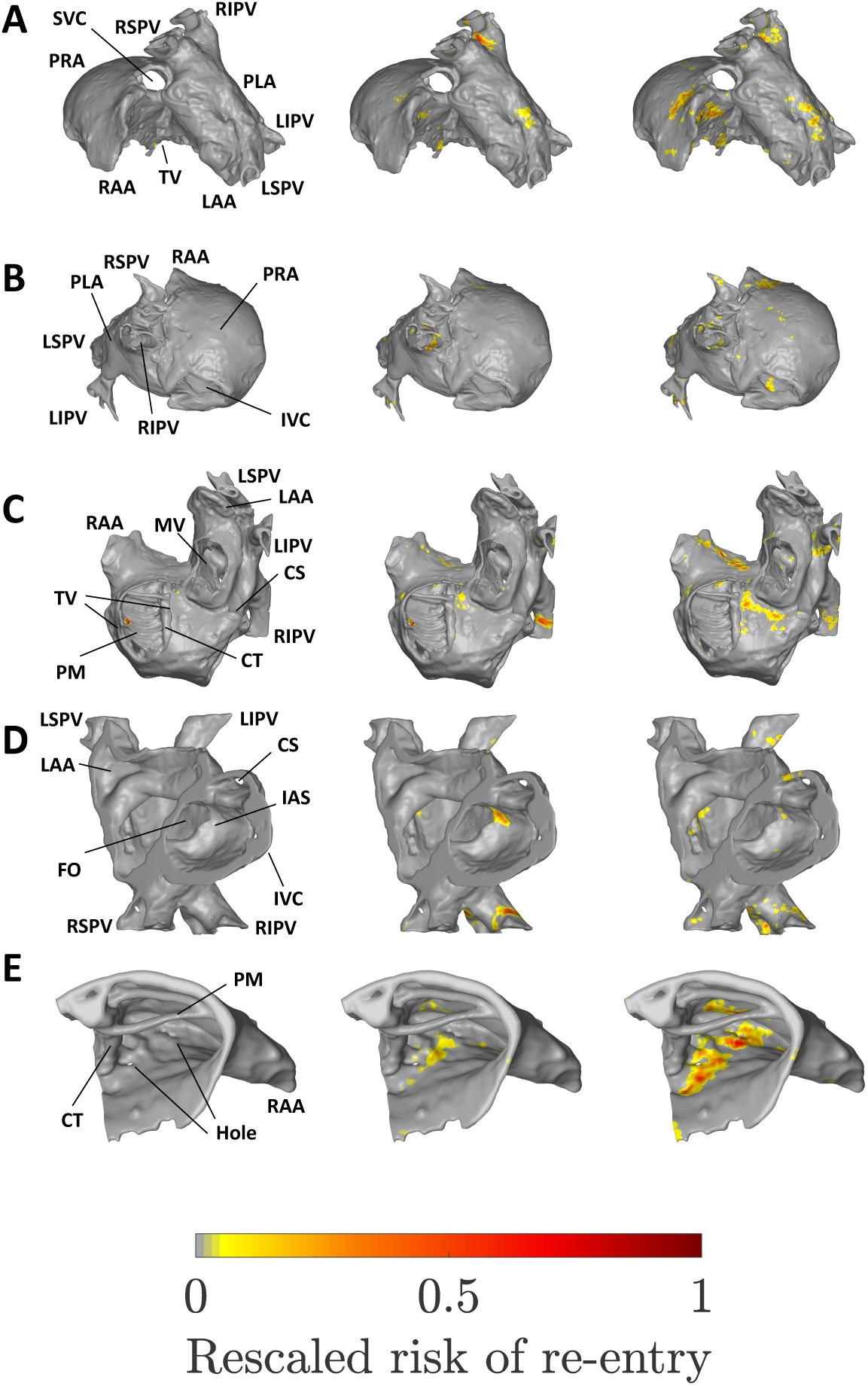
The micro-reentrant substrate for the human atria at low (left column), medium (middle) and high risk of micro-reentry (right). A: Superior view. B: Posterior view. C: Inferior view. D: Cut-through. E: Superior right atrial zoom. IVC/SVC: Inferior/ superior vena cava. PRA/PLA: Posterior right/ left atrium. RAA/LAA: Right/ left atrial appendage. TV/MV: Tricuspid/ mitral valve opening. CS: Coronary sinus. FO: Fossa ovalis. PM: Pectinate muscles. CT: Crista terminalis. IAS: Inter-atrial septum. RIPV/LIPV/RSPV/LSPV: Right/ left, inferior/ superior, pulmonary vein.

**Fig 6.**
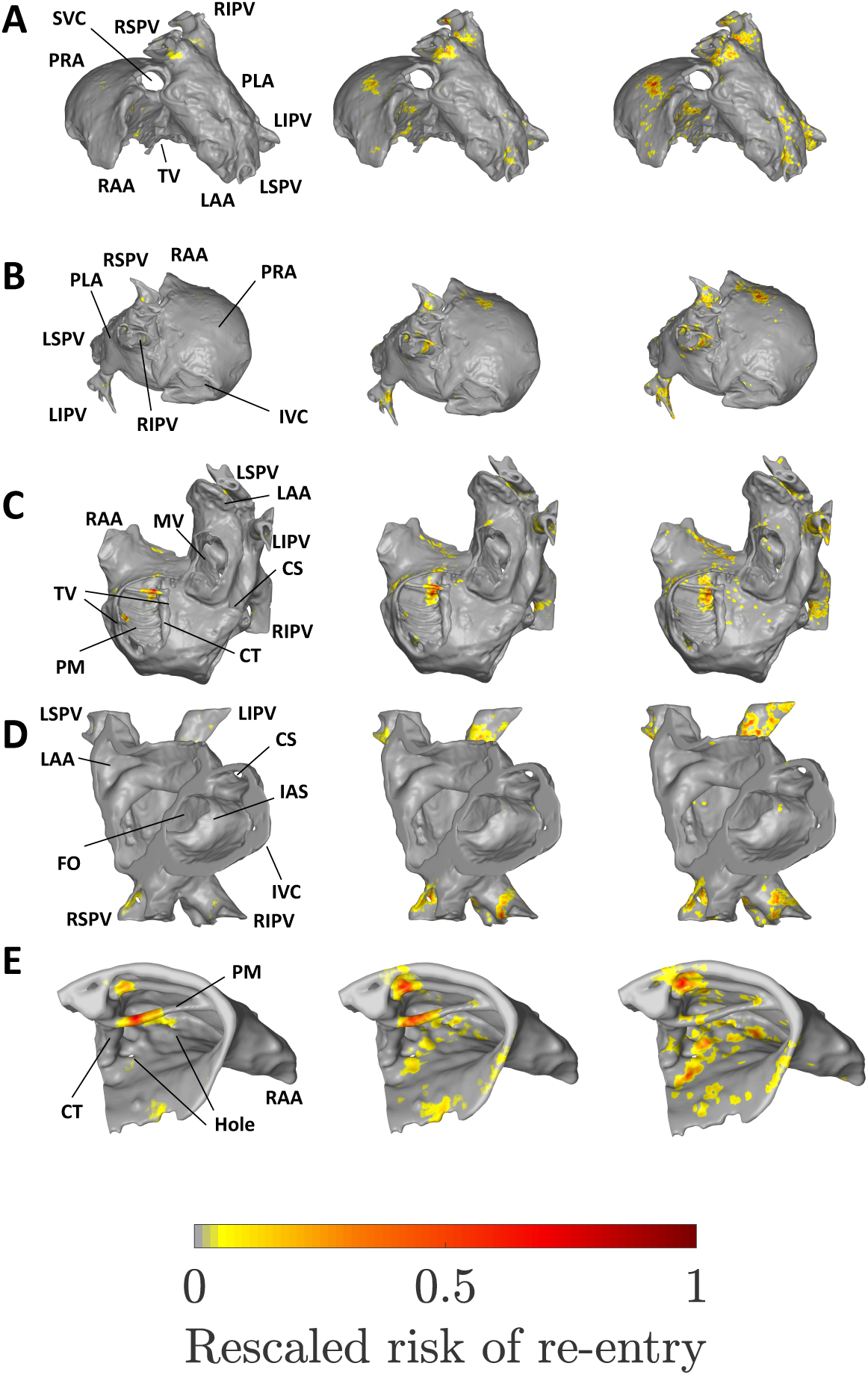
The micro-reentrant substrate for the human atria using the fibre-less null model at low (left column), medium (middle) and high risk of micro-reentry (right). A: Superior view. B: Posterior view. C: Inferior view. D: Cut-through. E: Superior right atrial zoom. See Fig. 5 for labels.

The reader should be conscious that the risk distributions shown do not represent the regions at risk for a single atrial tissue, but rather show the collation of the risk regions identified across 1000 randomly generated spatial networks. If a region is highlighted as at risk in the left column, this may be interpreted as an area which is susceptible to micro-reentry even if the fibrosis density is low (but non-zero). Regions identified in the middle and right columns are regions susceptible to micro-reentry at moderate and high fibrosis densities respectively. This illustrates a key result: different regions of the atria are susceptible to micro-reentry at different fibrosis densities. In particular, regions may have a characteristic fibrosis density range within which they are susceptible to micro-reentry; if the fibrosis density is too high or low these regions will not exhibit micro-reentry.

#### 4.1.1 Micro-reentrant substrate in spatial network with fibre structure

At low risk, micro-reentry is exclusively observed along a spatially isolated ridge of the pectinate muscles (PMs) in the right atrium, see Fig. 5. At medium risk, the pulmonary veins (PVs), in particular the right inferior pulmonary vein (RIPV), emerge as the dominant micro-reentrant substrate. Secondary substrates include the junction of the left LSPV and the PLA, and the inter-atrial septum (IAS). However, the hole in the IAS may play a role in it emerging as a risk region. An additional strip of low risk is observed in the superior right atrium, adjacent to the crista terminalis (CT), see Fig. 5(E). The strip in question lies between two small holes in the atrial dataset, with sub-millimetre diameters.

At high risk, the susceptible substrate is significantly more diffuse than at lower risk. However, there remain multiple regions with very low or zero risk, in particular the posterior right atrium (PRA), left atrial appendage (LAA) and inferior vena cava (IVC). The dominant risk substrate lies along the strip between two small holes in the superior right atrium. Secondary regions of high risk are maintained in the PVs, with risk migrating slightly further from the LSPV junction into the PLA. The opening of the the coronary sinus forms a new substrate which is not observed at higher coupling. The risk observed previously at the IAS and PMs is largely absent.

#### 4.1.2 Micro-reentrant substrate in spatial network without fibre structure

The risk substrate identified by the fibre-less null model, see Fig. 6, is qualitatively similar to the risk regions identified in the fibre model. Key risk regions including the superior right atrium and the PV sleeves are common to both models. Likewise, the PRA, LAA and IVC are not susceptible to micro-reentry in either model.

At low risk, the dominant risk substrate in the fibre-less null model is confined to two spatially isolated PM ridges, one of which exhibits risk in the fibre model. Additional diffuse risk is observed in the superior right atrium and in the right PV sleeves. At medium risk, the low risk substrate is retained, with additional risk in all four PVs and diffuse risk across the superior right atrium. These risk regions are consolidated at high risk, with a small reduction in risk observed along the spatially isolated PV ridges. The substrates observed for the fibre model along the coronary sinus (CS) opening and at the IAS are not observed in the fibre-less null model.

### 4.2 Clustering of micro-reentrant substrate

The observed micro-reentrant risk substrates for the fibre and fibre-less null models suggest that micro-reentry is spatially confined for both models at low risk, but that the substrate covers a wider area as the risk of micro-reentry increases. Figure 7 shows that the substrate is more clustered for the fibre model than for the fibre-less null model. One possible interpretation of this result is that if all micro-reentrant circuits are confined to a small number of clusters, these will be easy to destroy or isolate via catheter ablation. However, if circuits are widely distributed, they will be more difficult to destroy or isolate.

**Fig 7.**
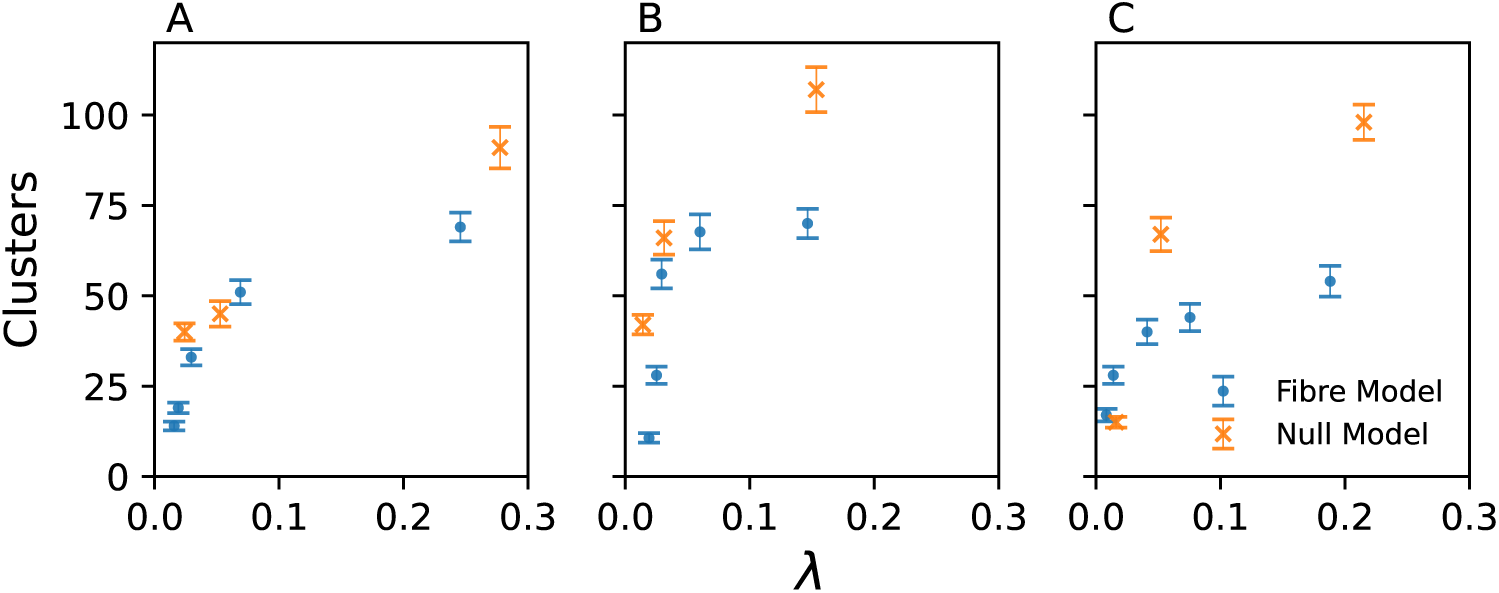
The number of independent spatial clusters identified as at risk of micro-anatomical reentry in the fibre model (blue points) and the fibre-less null model (orange crosses) for (A) the human atria, (B) the healthy sheep atria, and (C) the HF sheep atria, as a function of the risk parameter, *λ*. Each clustering value is calculated from 1000 sampled micro-reentrant circuits. Error bars from bootstrapping.

### 4.3 Occupied voxel fraction

Figure 8 shows the average occupied voxel fraction for the identified micro-reentrant substrates in the fibre and fibre-less null models. For all three atrial datasets, the figure demonstrates that at low risk, the micro-reentrant substrate is confined to regions with low OVF, significantly below the average value for each dataset. As risk increases (increasing *λ*), the mean OVF increases. This indicates that the bias to thin convex regions of the atria is reduced as edges are progressively removed in each spatial network. Framed in terms of fibrosis densities, this result illustrates that regions with low OVF are susceptible to micro-reentry at a lower fibrosis density than regions with larger OVF. These results are supported by statistical analysis in SM section 2.3.

**Fig 8.**
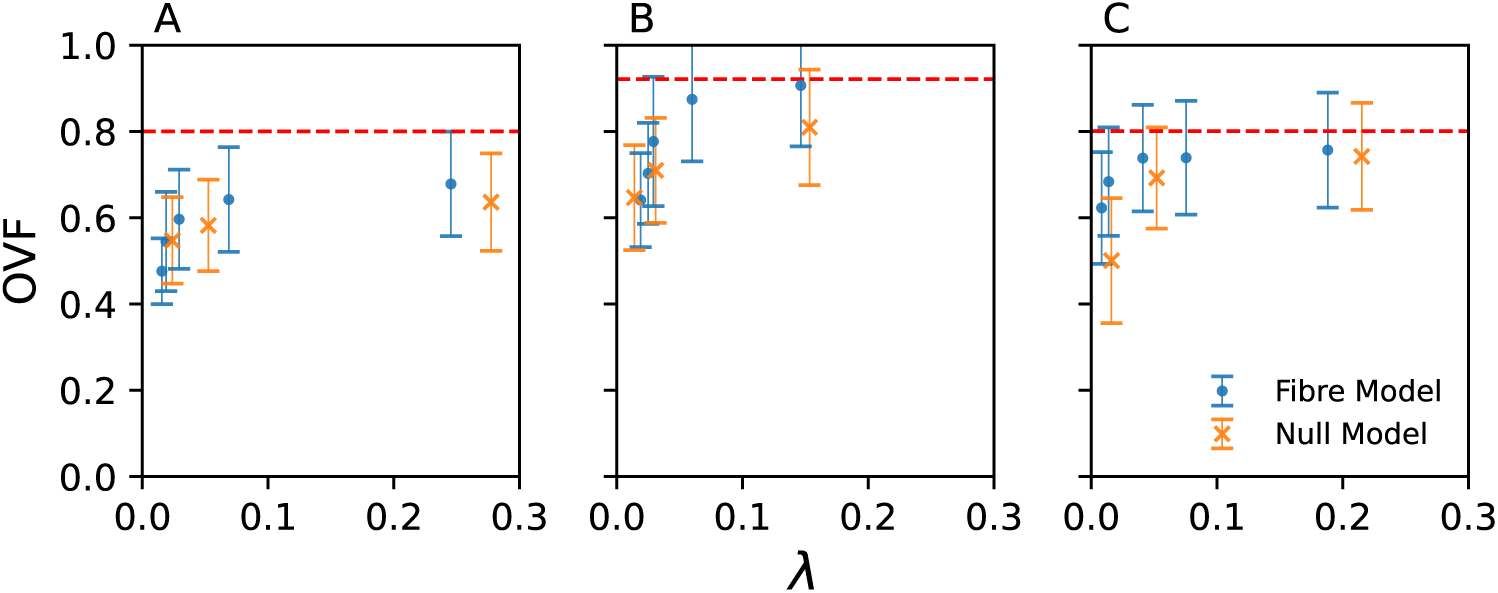
The occupied voxel fraction averaged over voxels where micro-reentry is detected in the fibre model (blue points) and the fibre-less null model (orange crosses) for (A) the human atria, (B) the healthy sheep atria, and (C) the HF sheep atria, as a function of the risk parameter, *λ*. The dashed line in each subfigure is the average value of the occupied voxel fraction for all the voxels in the atrial geometry.

### 4.4 Comparing fibre and fibre-less micro-reentrant substrates

Figure 9 shows an example of the micro-reentrant substrate for the fibre and fibre-less null models alongside their corresponding OVF and LCF values in the superior right atrium. The figure is for the human atria at high risk (right column in Figs. 5 and 6) where the bias to thin atrial regions is small but non-zero.

**Fig 9.**
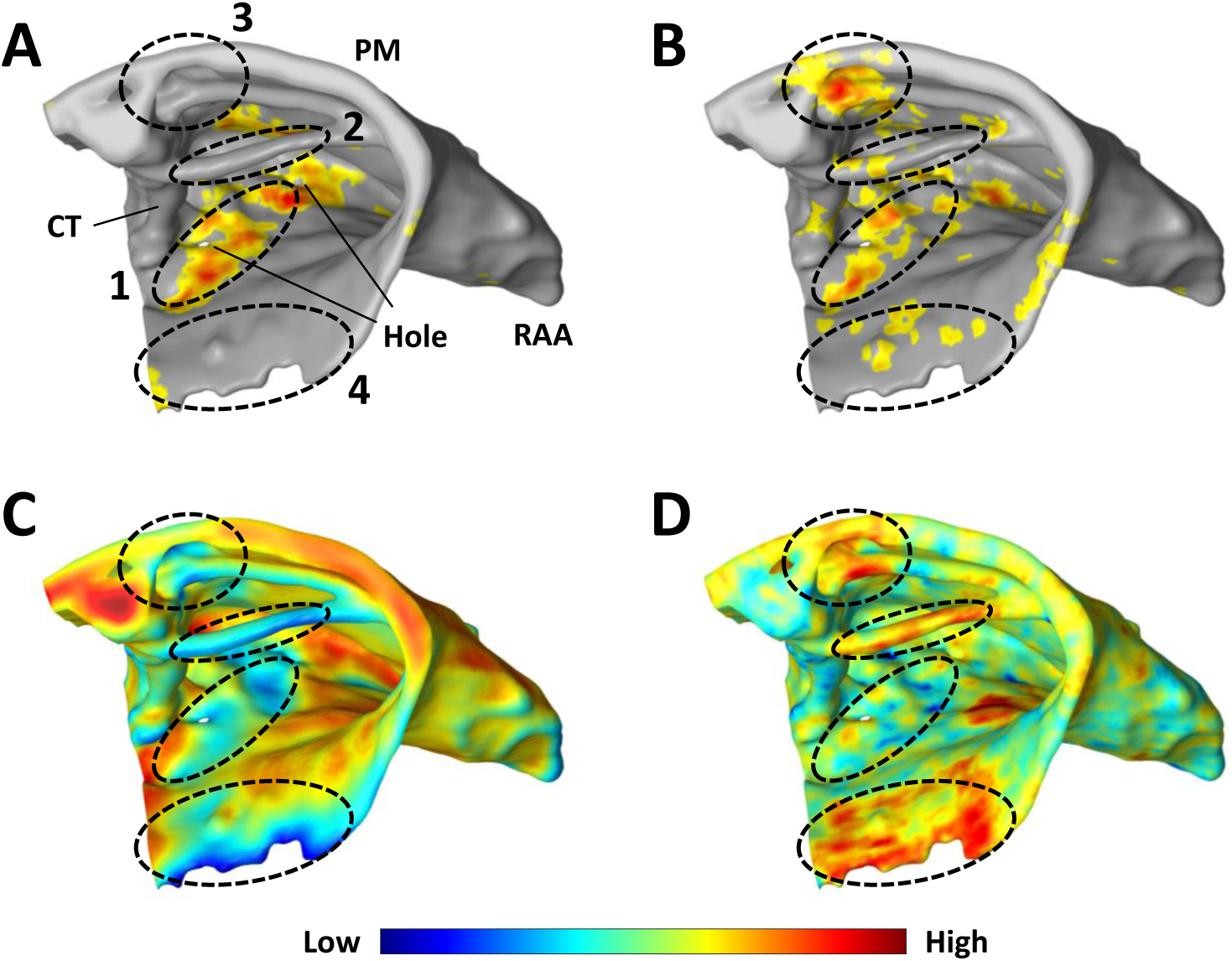
A view of the human superior right atrium illustrating factors influencing the spatial distribution of risk in (A) the fibre model, and (B) the fibre-less null model. (C) The occupied voxel fraction (OVF). (D) The longitudinal connection fraction in the fibre model (LCF). Regions (1–4) are referenced in the main text. Colourbar limits: OVF ∈ [0.3, 0.9], LCF ∈ [0.4, 0.8].

In the superior right atrium, the micro-reentrant substrate at high risk is concentrated along a strip between two small structural holes (region 1) for the fibre model, Fig. 9(A). This region exhibits moderate risk in the fibre-less null model, Fig. 9(B), with additional risk along the spatially isolated PM (region 2), the junction of the CT and a second PM ridge (region 3), and around the opening of the TV (region 4).

Region 2 exhibits low risk in the fibre model, despite being a low OVF region, but is the dominant risk region for the fibre-less null model when the rate of micro-reentry is low or medium, see Fig. 6(E) left and centre columns. This is because the strong longitudinal coupling in region 2 suppresses micro-reentry in the fibre model. This effect is absent in the null model where fibre structure is omitted. Region 2 is most susceptible to micro-reentry in the fibre-less null model at a low and moderate fibrosis density, but that the risk reduces at a very large fibrosis density, in contrast to some thicker regions of the atria. Regions 3 and 4 show very low or zero risk in the fibre-model.

Comparing the identified risk regions to the OVF values shown in Fig. 9(C), we note that regions 1–4 all exhibit low OVF values (blue on the colour scale). Of the four regions identified, we observe that the LCF values, Fig. 9(D), in region 1 are low (blue), indicating fibre misalignment, whereas strong longitudinal coupling is observed in regions 2–4. This suggests that the risk of micro-reentry is suppressed in the fibre model in regions with high LCF, but can be enhanced in regions with low LCF.

These patterns can be quantified by plotting the distribution of LCF and OVF values across all the voxels in the human atria. Figures 10(A) & (D) show the distribution where all voxels are equally weighted at (A) low and (D) high micro-reentrant risk. In Figs. 10(B) & (E) these distributions are weighted by the rescaled risk of micro-reentry observed in each voxel for the fibre model. The equivalent is shown in Figs. 10(C) & (F) where voxels are weighted by their risk in the fibre-less null model^1^.

**Fig 10.**
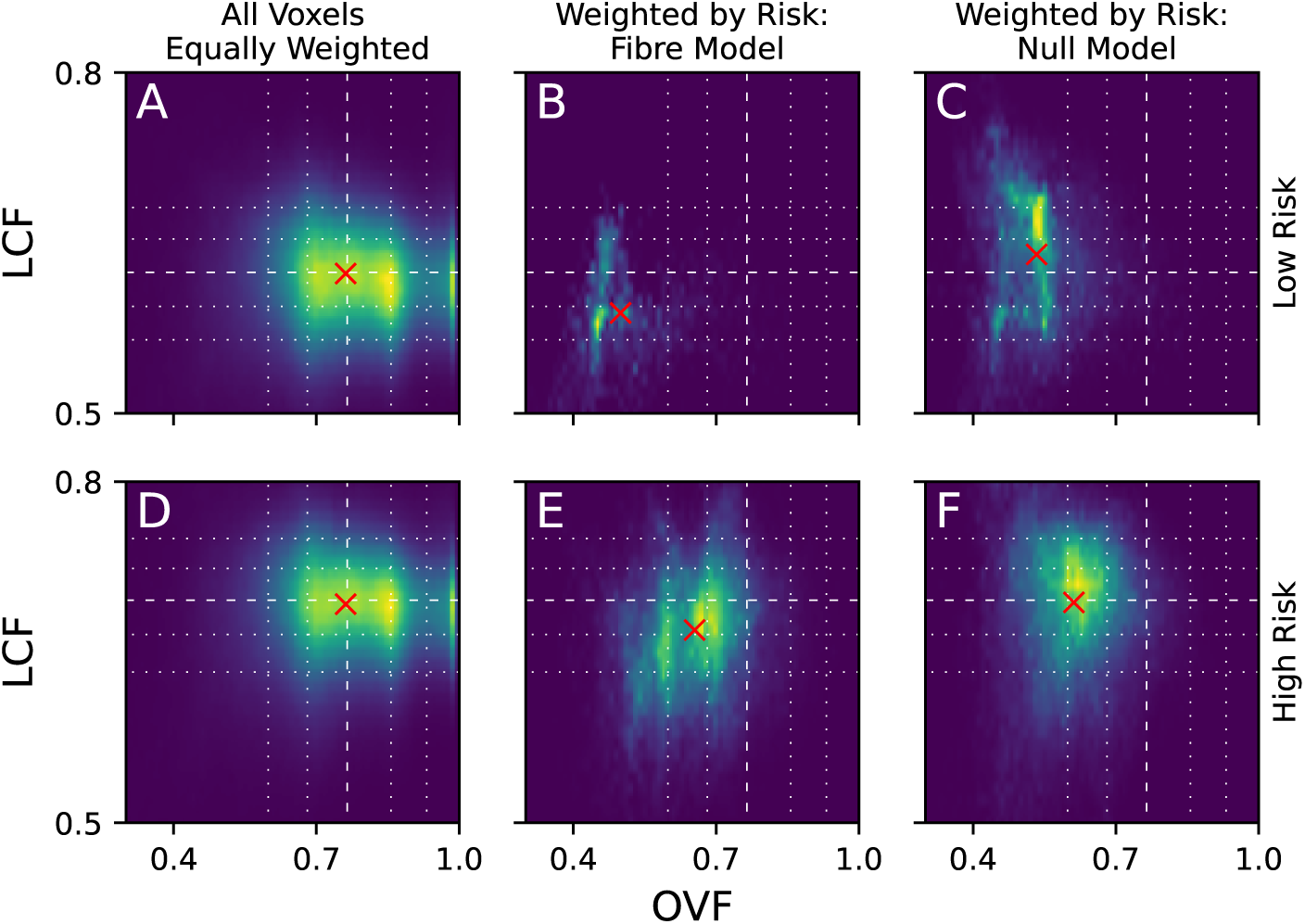
The distribution of LCF and OVF values in the human atria at low (A-C) and high (D-F) risk. (A) & (D) The distribution across all the voxels in the human atria. (B) & (E) The distribution weighted by the relative risk of reentry in each voxel for the fibre model. (C) & (F) The equivalent for the fibre-less null model. Note, the LCF is not defined for the fibre-less null model, but for illustrative purposes we include the LCF for the equivalent fibre model. Dashed lines: median. Dotted lines: 10^*th*^, 25^*th*^, 75^*th*^ and 90^*th*^ percentiles. Yellow: high voxel density. Blue: low voxel density. Red crosses: weighted mean.

If the distribution of micro-reentrant risk is unbiased by OVF or LCF, the distributions in Figs. 10(B), (C), (E) & (F) should follow the same patterns shown in Figs. 10(A) & (D). For the fibre model at low risk, Fig. 10(B), the voxel distribution is significantly skewed for both the OVF and the LCF with the weighted mean falling in the first OVF decile and the first LCF quartile. For the fibre-less null model at low risk, Fig. 10(C), the mean OVF value also falls in the first decile, but there is a small skew to LCF values larger than the median. This is an indirect effect and is explained by the strong longitudinal coupling along spatially isolated regions such as the PMs.

For both the fibre and fibre-less null models the risk density lies almost exclusively in the first OVF quartile, with negligible density at higher OVF values. In contrast, LCF values are clearly skewed in the fibre model, although some risk density remains at larger LCF values. This implies that at low risk, the OVF is the dominant factor in determining the micro-reentrant risk substrate, with fibre structure playing an important secondary role.

As the risk of micro-reentry increases, we observe that the bias to low OVF values remains, but is slightly reduced relative to the low risk cases, see Figs. 10(C) & (F). However, the risk of micro-reentry is still completely suppressed at the largest OVF values. For the fibre model, a small bias towards reduced LCF is observed. In contrast, the fibre-less null model shows minimal bias in the distribution of LCF values. Statistical arguments supporting the above discussion are provided in SM section 2.3.

## 5 Discussion

In this paper, we introduce a novel computational approach for analysing how the structural substrate for micro-anatomical reentry may develop as the atria accumulate diffuse interstitial fibrosis. The technique combines spatial networks with fibre tractography, and assumes that interstitial fibrosis accumulation can be modelled as the decoupling of a spatial network, a concept closely related to spatial percolation [11].

Many studies investigate the electro-anatomical basis for micro-anatomical reentry. Broadly speaking, such studies fall into three categories. In the first, the substrate can be identified directly, given sufficiently high resolution imaging, by observing the presence of a micro-anatomical driver. Examples include [7, 8], identifying several key factors in the formation of the reentrant substrate. Of particular note are the orientation of muscle fibres, the structure, thickness, and thickness gradients of the atria, and the accumulation of fibrosis, particularly interstitial fibrosis [27].

Attempting to quantify these factors, the authors in [8] applied optical mapping to explanted human atria to identify how specific values of wall thickness and fibrosis burden correlate to the location of the known micro-reentrant substrate [8]. Driver regions were found to correlate well with areas of 20-30% wall thickness and 20-30% fibrotic burden, notably the junction of the RIPV and PLA, and between the CT and fibrotically insulated PMs. Fibre misalignment was also implicated, specifically between the CT and the PMs which also exhibit abrupt changes in the local wall thickness. However, these studies took a largely static view of the micro-reentrant substrate, not considering how the substrate may change under electrical or structural remodelling.

In the second category, the substrate for micro-reentry has been established in the atria, but is hidden due to the lack of a clearly observable driver. One recent study has addressed this problem by noting that the visible micro-reentrant substrate varies strongly with variable atrial refractoriness [28]. Hence, the authors were able to demonstrate that the hidden micro-reentrant substrate could be unmasked and stabilised by shortening atrial refractoriness with adenosine. Such changes, induced pharmacologically, are analogous to some of the electro-mechanical changes that may be expected over time from atrial remodelling [4]. However, a robust framework for predicting the future micro-reentrant substrate is still lacking.

No study is yet to address the third category which is to predict how the substrate for micro-reentry will develop in the future on a patient specific basis, a problem directly related to the increasing need for arrhythmic risk stratification [29].

In this proof of concept – which we acknowledge is far from clinical applicability – we have attempted to take a step towards addressing this and ask how the substrate for micro-reentry may develop over time. Specifically, we ask how different parts of the atria are susceptible to micro-reentry at different characteristic fibrosis ranges; some regions of the atria, particularly those where the atrial walls are thin and there is significant fibre misalignment, may be susceptible to micro-reentry at a low fibrotic density, whereas other regions may require higher fibrosis densities before micro-reentry can be induced. Such an approach is currently only possible in-silico, primarily due to ongoing challenges with accurate in-vivo atrial imaging [30].

Our results implicate the role of wall thickness and the misalignment of fibres, suggesting that thin atrial walls and reduced longitudinal fibre coupling both enhance the probability that a region is susceptible to micro-reentry at lower fibrosis densities, supporting the findings in [8]. However, our study suggests that the dependence on these factors evolves as the density of interstitial fibrosis grows in the atria. In particular, our study suggests that the spatial spread of the micro-reentrant substrate increases dramatically with small reductions in the spatial network coupling, and indicates that the bias to thin atrial regions with complex fibre morphology is reduced as the micro-reentrant substrate becomes more spatially diverse. Many of the specific regions highlighted as risk substrates for micro-reentry in [8] and [28], such as PM ridges and the superior right atrium, naturally emerge as risk substrates using our method, most likely due to their position in thin atrial regions.

One explanation for the importance of wall thickness, fibrosis density, and local wall curvature is that driver regions emerge from percolation-like dynamics where the critical fibrosis density is dependent on the thickness of the structure studied [31, 32]. This in turn may result in micro-anatomical circuits anchoring to the atrial surfaces in paroxysmal AF but distributing across the atrial wall in persistent AF [33]. Aside from micro-reentry, modelling fibrosis using percolation-style distributions is known to perform particularly well when simulating patterns of AF maintenance [34], and has been used to explain complex fractionated electrograms and reentries in fibrotic border zones [35, 36].

In the wider AF literature, a number of computational studies consider the role of atrial structure on AF dynamics. In most cases, these studies investigate how structural factors effect electrical wavefront dynamics by solving the mono- or bidomain equation coupled with a suitable atrial cell model [32, 37–42]. However, even the most advanced patient-specific modelling methodologies suffer from the continued struggle to extract precise fibre orientation data and high resolution fibrosis profiles (particularly diffuse interstitial fibrosis) in-vivo [6].

Until these issues are resolved, computational modelling must continue to be used as a platform for hypothesis testing, studying the dynamics of AF initiation and maintenance with a variety of methods and across scales. In particular, the clearest insights may be found from models which contrast the role of a specific electro-anatomical feature with null models in the absence of that feature. Examples include models with and without realistic tissue geometries [37], with and without patient-specific fibrosis [43], isotropic vs. anisotropic fibre structures [40], and continuous vs. discrete modelling modalities [24]. We hope that the techniques introduced in this paper can add to the range of techniques employed for hypothesis testing in cardiac electrophysiology.

### 5.1 Limitations

The aims of this study are highly focused, discussing the structural basis for micro-anatomical reentry only. Many other factors may effect the probability that a micro-reentrant substrate forms in a given region of the atria, including other forms of fibrosis and ion channel remodelling. Our approach does not consider these factors and cannot exclude their importance to micro-anatomical reentry.

The key technical limitations of our work come under two main categories: (1) Imaging related limitations and (2) limitations arising from specific modelling choices in the construction of the spatial networks. These are discussed fully in SM section 1.9. In our view, the most important limitation relates to the datasets used for analysis. In particular, the human fibre dataset has undergone extensive smoothing, lacking realistic local heterogeneity in the fibre structure; the observed heterogeneous regions are likely artefacts from the synthetic fibre generation method. Similarly, the atrial geometry is heavily smoothed, lacking fine structural detail in regions such as the RAA and LAA, but with numerous structural holes across the geometry. Any future work must ensure that predictions are not simply an artefact of the data acquisition and generation process.

Although using high resolution fibre maps like those in [15, 16] may avoid these issues, acquiring such data is not feasible in-vivo, remaining a major ongoing research challenge [44, 45].

## 6 Conclusion

We have introduced a simple, proof of concept framework which attempts to study how the substrate for micro-anatomical reentry develops from the accumulation of interstitial fibrosis. The method, which is based on the application of percolation to spatial networks, suggests that the micro-reentrant substrate is critically dependent of local tissue geometry and areas of fibre misalignment. We suggest that the dependence on these factors is complex, continuously evolving with the absolute level of micro-reentrant risk.

## 8 Data accessibility

The three atrial fibre datasets are available for download from doi.org/10.5281/zenodo.5500725

The code used to generate the spatial networks from the atrial imagine data is available at github.com/JamesAlecColeman/reentrySpatialNetworks.

Code and subsidiary data used to process the spatial network data and generate our results is available at doi.org/10.5281/zenodo.5500003

## 9 Acknowledgements

MF is grateful to Rebecca Joakim for drawing the schematics in Figures 1 and 2, and thanks Hardik Rajpal for help with the statistical analysis. MF is also grateful to Tim S Evans for a number of useful discussion throughout the project.

MF acknowledges a PhD studentship from the Engineering & Physical Sciences Research Council (GB) via grant number EP/N509486/1. AC acknowledges a PhD studentship from the Engineering & Physical Sciences Research Council (GB) via grant number EP/L015129/1. NSP and FSN are supported by the Imperial Centre for Cardiac Engineering and by the British Heart Foundation through programme grant number RG/16/3/32175. FSN is supported by the National Institute for Health Research and the Imperial Biomedical Research Centre. The funders had no role in study design, data collection and analysis, decision to publish, or preparation of the manuscript.

## 10 Author Contributions

MF and KC proposed the project and developed the key ideas. MF, JC, SD and KC developed the final version of the original methods. JC, SD, DJH and LT implemented the methods, supervised by MF, AC and KC. Datasets and technical expertise were provided by BT and JZ. Medical expertise were provided by FSN, SA, NSP and JZ. The manuscript was written by MF and JC, with comments and edits provided by KC, JZ, SA and FSN.

## Footnotes

1 Although the LCF is not defined for the fibre-less null model, it is illustrative to label null model voxels according to the corresponding fibre model LCF values.

